# Identifying robust strategies for assisted migration in a competitive stochastic metacommunity

**DOI:** 10.1101/811497

**Authors:** Gregory A. Backus, Marissa L. Baskett

**Affiliations:** University of California, Davis

## Abstract

Assisted migration is the translocation of species beyond their historical range to locations that are expected to be more suitable under future climate change. However, a relocated population might fail to establish within its donor community if there is high uncertainty in decision making, climate, and interactions with the recipient ecological community. To quantify the benefit to persistence and risk of establishment failure of assisted migration under different management scenarios, we built a stochastic metacommunity model to simulate several species reproducing, dispersing, and competing on a temperature gradient as temperature increases over time. Without assisted migration, the species in our model were vulnerable to climate change if they had low population sizes, short dispersal, and strong poleword competition. When relocating species that exemplified these traits, assisted migration increased the long-term persistence of the species most when relocating a fraction of the donor population, even if the remaining population was very small or rapidly declining. This suggests that leaving behind a fraction of the population could be a robust approach, allowing managers to repeat assisted migration in case they move the species at the wrong place and wrong time, especially when it is difficult to identify a species’ optimal climate. We found that assisted migration was most beneficial to species with low dispersal ability and least beneficial to species with narrow thermal tolerances, for which assisted migration increased extinction risk on average. Lastly, while relocation did not affect the persistence of non-target species in our simple competitive model, researchers will need to consider a more complete set of community interactions to comprehensively understand invasion potential.

## Introduction

Global biodiversity is expected to decline with projected climate change (Urban 2015). Among the species at risk of extinction are those with limited dispersal, narrow ranges, narrow climate tolerance, and low population sizes (Pearson 2006; Tewksbury et al. 2008). Competition and other community interactions could increase extinction risk, as negative interactions can limit the dispersal of species that might be otherwise adequate dispersers (Gilman et al. 2010; Urban et al. 2012). Many of these climate-threatened species face a high likelihood of extinction without human intervention, prompting scientists and managers to consider a variety of novel approaches to conservation (Heller & Zavaleta 2009). Among these approaches is assisted migration (AM), in which managers relocate individuals from a threatened population to a location outside their historical range that is expected to be more suitable under future climates (McLachlan et al. 2007). By allowing these species to reach favorable climates in densities that they would not be able to reach on their own, AM could increase the likelihood of persistence for climate-threatened species or protect species with high ecological and socioeconomic value in ways that traditional conservation strategies might not (McLachlan et al. 2007; Heller & Zavaleta 2009; Lawler & Olden 2011; Schwartz et al. 2012).

Moving a species into a novel ecosystem could also incur many risks (Ricciardi & Simberloff 2009; Hewitt et al. 2011). Ecological risks include the risk that a relocated species becomes invasive within its recipient community (Mueller & Hellmann 2008), the risk of spreading pathogens and parasites (Simler et al. 2018), demographic costs of increased elevated stress on individuals when relocating individuals (Dickens et al. 2010), and the loss of genetic diversity when creating small founding populations and donor populations (Kekkonen & Brommer 2015). Demographic and genetic risks might contribute to the risk of a species failing to establish after relocation (Chauvenet et al. 2013; Plein et al. 2016) as seen with the low-to-intermediate success of previous conservation translocations across a wide range of taxa (Fischer & Lindenmayer 2000; Godefroid et al. 2011). Establishment failure could accelerate the negative impacts on climate-threatened species by reducing the population size and genetic diversity while diverting the limited resources available to conservation (McDonald-Madden et al. 2008; Hewitt et al. 2011). This risk also depends, in part, on uncertainties that lead managers to relocate a species into the wrong place at the wrong time, especially if there are narrow conditions under which a species can persist. One source of uncertainty that leads to translocation failures is environmental stochasticity (Wolf et al. 1996), which will likely increase with climate change (Vasseur et al. 2014). Additional uncertainty stems from the difficulty in quantifying and differentiating between the abiotic and biotic drivers of species’ ranges (Case et al. 2005), which could become increasingly uncertain with climate change (Boiffin et al. 2017). Given these uncertainties, a key management challenge is developing robust approaches over a range of conditions (McDonald-Madden et al. 2008) for the array of decisions involved in AM. This involves evaluating which species are vulnerable to climate-threatened extinction, which species will likely benefit from AM, when and where to move a species, and how many individuals to move (McDonald-Madden et al. 2011; Rout et al. 2013).

Despite a lack of consensus among the scientific community and the public about the benefits and risks of AM (Hewitt et al. 2011; Javeline et al. 2015; St-Laurent et al. 2018), several species are already being relocated (Seddon et al. 2015). Scientific guidance for AM endeavors is available from existing AM decision-making frameworks, which typically focus on optimizing a species’ persistence under climate change using single-species models (McDonald-Madden et al. 2008; Rout et al. 2013; Kling et al. 2016). Extending these to a multispecies framework is a crucial next step to account for the species interactions that give rise to the risks of invasiveness and uncertainty in the drivers of species range.

In this paper, we quantify the benefit to target species’ persistence, risk of failed translocation, and risk of non-target species’ response to AM given competitive interactions, multiple sources of uncertainty, and an array of management decisions. We built a stochastic metacommunity model to simulate competing species undergoing climate change to estimate which species were vulnerable to extinction, which species were likely to benefit from AM, and what fraction of the population to relocate. Though the dynamics of interacting communities that give rise to the risk of invasiveness are often complex, involving many direct and indirect trophic interactions (Holt 1984; Chesson & Kuang 2008), we focus on competition because of its role in driving species’ range limits, rooted in long-standing ecological theory (Connell 1972) and established as the most extensively studied and best-supported species interaction for driving range limits (Sexton et al 2009). Due to this role in affecting species’ ranges generally, competition has a significant potential to affect range shifts and persistence under climate change (Urban et al. 2012; Ettinger & HilleRisLambers 2013), the dynamic central to the AM decisions on which we focus. Another potential source of uncertainty comes from abiotic drivers of species’ ranges. Because managers will have limited knowledge of a species’ optimal climate (a reducible uncertainty), we simulated relocation with uncertainty in estimating of species’ thermal optima. By repeating these simulations under different levels of environmental stochasticity (an irreducible uncertainty), we identified characteristics of successful AM approaches that were robust over a wide variety of uncertainty scenarios.

## Methods

### Model overview

To compare assisted migration (AM) strategies, we modeled metacommunity dynamics of multiple species competing on a one-dimensional linear temperature gradient subjected to climate change, analogous to a previous model by Urban et al. (2012) with environmental stochasticity. For simplicity, all species in this metacommunity are annuals competing over the same resources at the same trophic level. The model cycles through reproduction, dispersal, and competition, all with demographic stochasticity, over each time step (Fig. 1). Each species *i* has a unique dispersal distance (*γ*_*i*_), thermal optimum (*z*_*i*_), thermal tolerance breadth (*σ*_*i*_), and a reproductive strength parameter (*r*_*i*_) that scales the birth rate to create a specialist/generalist trade-off (Levins 1968). We simulate AM by selecting one target species and relocating a fraction of its total population toward its thermal optimum each time the population falls below a threshold population size. We compared outcomes when relocating different target species with different fractions of the population into different locations.

**Figure 1:**
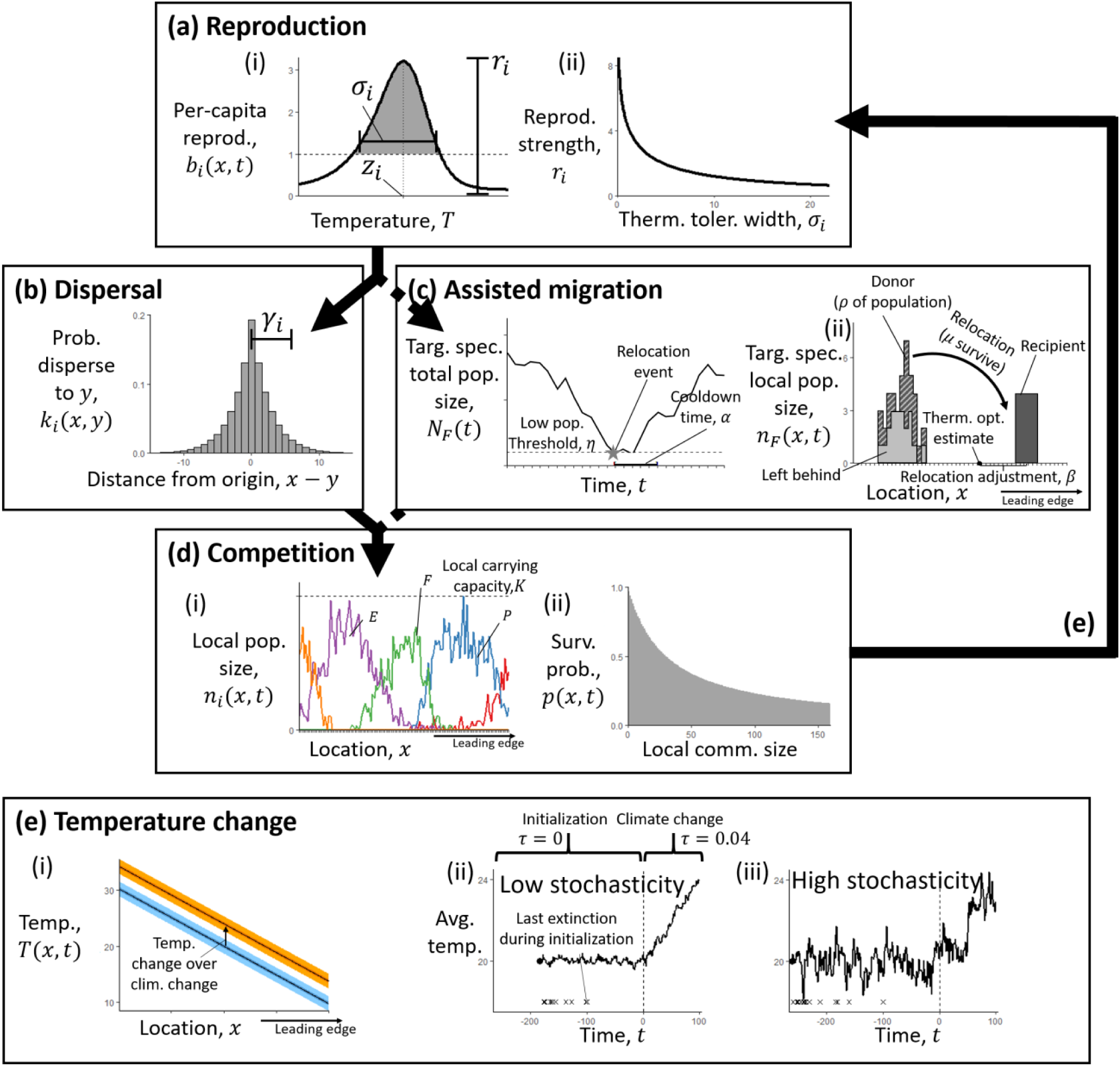
During each time step of the model, all extant species cycle through (a) reproduction, (b) dispersal, and (d) competition before (e) the temperature changes and the next time step continues. The target species also experiences (c) assisted migration during certain time steps. (a.i) Per capita reproductive output *b*_*i*_(*T*(*x, t*)) is skew-normal, dependent on temperature *T*(*x, t*). This function is shaped by species’ thermal optimum *z*_*i*_ and thermal tolerance breadth *σ*_*i*_. (a.ii) Reproductive strength *r*_*i*_ scales the total reproductive output so that species with narrow *σ*_*i*_ (specialists) have higher reproduction and species with broad *σ*_*i*_ (generalists) have lower reproduction. (b) The dispersal kernel is a long-tailed “double geometic” distribution with a mean dispersal distance *γ*_*i*_. (c.i) Relocation occurs once the total population of target species *F* falls below a threshold *η*. To avoid repetition while *F* recovers, no relocations occur during a cool-down period following relocation *α*. (c.ii) A fraction *ρ* of population *F* is removed from its original distribution and moved to a new location (only *μ* survive) *β* patches beyond the leading edge. Remaining individuals disperse naturally. (d.i) All species compete over limited space, where each patch has a carrying capacity *K*. Here each line represents a different species. (d.ii) In each patch, individual survival probability *p*(*x, t*) decreases as the total community size increases. (e) Temperature changes stochastically over time. (e.i) Mean temperature decreases linearly with space. Over time, between *t* = 0 (lower line) and *t* = 100 (upper line), the temperature increases by 4°C. (e.ii-iii) Temperature variation over time depends on level of environmental stochasticity. Both examples have the same autocorrelation (*κ*), but (ii) has a higher standard deviation (*ψ*). The vertical dashed line designates when the model changes from the initialization phase (average temperature change (*τ* = 0)) to the climate change phase (*τ* = 0.04). Climate change only occurs after a relatively stable metacommunity has been assembled, after 100 time steps have passed with no extinctions.

We analyze the potential benefit of AM based on the target species’ persistence likelihood throughout all locations. Persistence likelihood also indicates the risk of translocation failure due to uncertainty and stochasticity. We measure the risk of invasiveness from competitive interactions based on the persistence likelihood of non-target species as well as the overall gamma diversity. Though this metric only represents an extreme outcome on the continuum of possible invasive impacts on community structure and ecosystem function (Blackburn et al. 2014), other metrics would require investigation beyond competitive interactions within a trophic level.

### Population dynamics

Each species *i* has a local population size of *n*_*i*_(*x, t*) individuals in discrete patch *x* and a total metapopulation size over all space *X* of *N*_*i*_(*t*) = ∑_*x*∈*X*_ *n*_*i*_(*x, t*) at discrete time *t*. First, all individuals reproduce (Fig. 1a) with a reproductive output *b*_*i*_(*T*(*x, t*)) that depends on local temperature *T*(*x, t*). Temperature-dependence is skew-normal, given skewness constant *λ*, and highest values are around the thermal optimum *z*_*i*_ with a sharp decrease above *z*_*i*_ (Norberg 2004). Thermal tolerance breadth *σ*_*i*_ and reproductive strength *r*_*i*_ determine the breadth and height of the temperature-dependence. Altogether, *b*_*i*_(*T*(*x, t*)) is

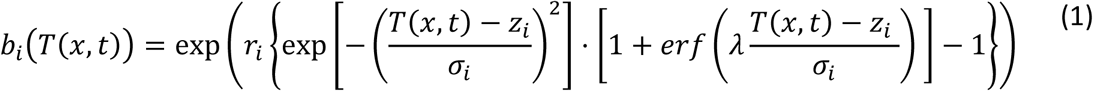

(following Urban et al. 2012). To incorporate demographic stochasticity, the number of propagules produced by species *i* in patch *x* is a Poisson random variable with mean equal to the reproductive output, 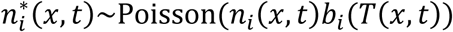 ∼Poisson(*n*_*i*_(*x, t*)*b*_*i*_(*T*(*x, t*)) (Melbourne & Hastings 2008).

Next, each propagule disperses from its origin (Fig. 1b). We adapted the Laplace dispersal kernel to a discrete-space analog (Appendix S1). We define *γ*_*i*_ as the mean absolute distance (in patches) that species *i* moves from its origin and let kernel parameter 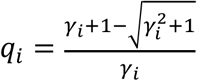. Thus, the probability of a propagule from patch *x* moving to patch *y* is

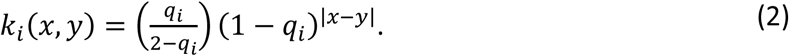

All propagules of species *i* disperse from patch *x* throughout all space *X* with the random vector 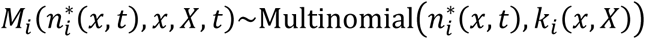, and the vector of local populations after dispersal is 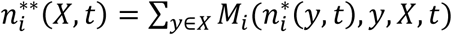.

Lastly, dispersed propagules compete within each patch *x* given community-wide carrying capacity *K* (Fig. 1d). We assumed a variation on lottery competition (Sale 1978; Chesson & Warner 1981), where each individual has an equal probability of surviving,

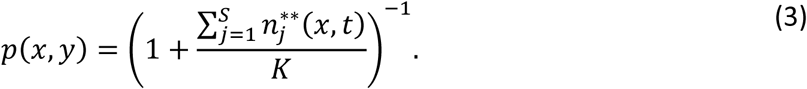

The total number of propagules of species *i* in patch *x* that survive after competition is a binomial random variable 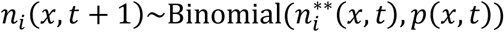 (Melbourne & Hastings 2008). Therefore, the overall quality of a location, in terms of how it drives production and survival, depends on the constant carrying capacity *K* and match between the local environment and species thermal tolerance.

### Spatial structure

Metacommunity dynamics occur across a one-dimensional, linear temperature gradient of *L* patches (Fig. 1e), representing gradual latitudinal or sharp elevational change (Urban et al. 2012). We remove propagules that disperse outside of the spatial gradient. Because these absorbing boundary conditions could bias our analyses, we disregard the first 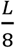 patches on the poleward edge and the last 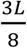 patches on the equatorward edge when measuring species and community metrics.

Temperature changes each time step by mean *τ* with autocorrelation *κ* and standard deviation *ψ* around white noise *ω*(*t*). The stochastic component of yearly temperature change is 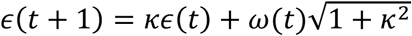, with the square root term to remove the effect of autocorrelation on the variance (Wichmann et al. 2005). Altogether, the temperature in patch *x* changes over time as

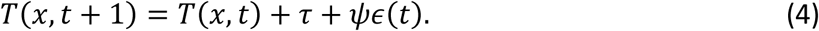

### Assisted migration

AM focuses on a single target species (Fig. 1c), species *F*. We relocate species *F* if its total metapopulation size *N*_*F*_(*x, t*) is below a threshold *η* at the beginning of a time step. Though population size would realistically need to be estimated, we assume perfect knowledge of population size to focus on uncertainty in the drivers of a species’ range. To avoid repeating AM before species *F* recovers, we only relocate if we did not previously relocate within the last *α* time steps. After reproduction, we select a fraction of propagules *ρ* for AM, randomly chosen from throughout the species’ range, while the remaining propagules are left behind to disperse naturally (the donor population). From the propagules chosen for AM, only a proportion *μ* survive relocation, and those are relocated uniformly among five connected patches around a patch *β* spaces poleward of the patch that most closely matches the species’ estimated thermal optimum (the recipient population).

We considered three methods of estimating the thermal optimum of species *F*. The perfect knowledge estimate is the exact value of the true thermal optimum *z*_*F*_. The realized niche estimate is the temperature in the median patch of the target species’ distribution at *t* = 0. The fundamental niche estimate measures species’ limits without competition by simulating 100 time steps with *τ* = 0°C/year and only species *F*. This estimate is the temperature in the median patch of the resulting distribution.

### Parameterization and implementation

Simulations occurred on a temperature gradient with *L* = 512 patches, where initial temperatures linearly varied over space from 9.78°C to 30.22°C. We considered two types of environments, defined by the degree of stochasticity. Low-stochasticity environments had an annual temporal standard deviation of *ψ* = 0.1639°C, equal to the standard deviation of mean combined global land-surface air and sea-surface water temperature anomalies from 1880 to 1979 (GISTEMP Team 2019; Lenssen et al. 2019), and high-stochasticity environments had four times that amount. Both had an annual temporal autocorrelation of *κ* = 0.767, also from temperature anomalies from 1880 to 1979. We used skewness constant *λ* = −2.7 (Urban et al. 2012) and carrying capacity *K* = 30 individuals. We chose this parameter space, including low values for *K*, so that extinctions do occur for some species and AM is a relevant management consideration (Appendix S3).

Before simulating climate change, we initialized the model to assemble a metacommunity with multidecadal coexistence under background environmental stochasticity. First, we generated a pool of *S* = 32 species, each with uniquely randomized dispersal distances *γ*_*i*_, thermal optima *z*_*i*_, and thermal tolerance breadths *σ*_*i*_, (based on Urban et al. 2012) (Table 1). We numerically derived the reproductive strength *r*_*i*_ from *σ*_*i*_, such that each species had the same overall reproductive potential *B* = 5 when integrating over temperature, emulating a jack-of-all-trades-master-of-none trade-off (Levins 1968). We placed 25 individuals from each species into five adjacent patches that most closely matched each species’ thermal optimum and iterated through the model with mean temperature change *τ* = 0 °C/year until 100 time steps passed without extinction. The remaining communities set initial conditions for subsequent climate change simulations (Appendix S4).

**Table 1:** Definitions of the symbols used in the model.

| Parameter | Symbol | Values | Units |
| --- | --- | --- | --- |
| Total species | $S$ | 32 | species |
| Dispersal distance of species $i$ | $\gamma_i$ | Lognormal; mean=2.5, st. dev.=2.5 | patches |
| Thermal optimum of species $i$ | $z_i$ | Uniform; 9.78 to 30.22 | °C |
| Thermal tolerance breadth of species $i$ | $\sigma_i$ | Lognormal; mean=5, st. dev.=5 | °C |
| Reproductive strength of species $i$ | $r_i$ | Derived from $\sigma_i$ | - |
| Skewness constant | $\lambda$ | -2.7 | - |
| Fraction of population relocated | $\rho$ | 0, 0.05, 0.1, ..., 1 | - |
| Assisted migration survival probability | $\mu$ | 0.8 | - |
| Low population threshold | $\eta$ | 42 | individuals |
| Cooldown time between relocations | $\alpha$ | 5 | years |
| Relocation adjustment (relative to optimum) | $\beta$ | 10 | patches |
| Total patches | $L$ | 512 | patches |
| Patch carrying capacity | $K$ | 30 | Individuals |
| Mean annual temperature change | $\tau$ | 0.04 | °C/year |
| Annual temporal autocorrelation | $\kappa$ | 0.767 | - |
| Annual temporal standard deviation | $\psi$ | low=0.1639, high=0.6556 | °C |
| Initial total population size of species $i$ | $N_i(0)$ | - | individuals |
| Difference in thermal optimum with species $i$ | $z_{\text{diff},i}$ | - | °C |
| Inverse Simpson's index of region $W$ | $D_W$ | - | - |
| Measured temperature change | $c_T$ | - | °C |
| Measured SD in temperature | $s_T$ | - | °C |
| Deviation in thermal optimum estimate | $z_{\text{est,dev}}$ | - | °C |

We modeled metacommunity dynamics under climate change with AM to test the success of a suite of potential relocation decisions. We initialized 10,000 metacommunities under both low and high stochasticity before iterating through 100 time steps with mean annual temperature change *τ* = 0.04°C/year to reflect projected temperature changes under RCP8.5 (Urban et al. 2012, IPCC 2014). For AM, we chose several target species that could be considered vulnerable to climate change, including: the species with the shortest dispersal, the species with the narrowest thermal tolerance breadth, the species with the closest poleward neighbor, the species with the lowest initial population size, and a randomly selected species. All target species were initially extant within an interior region of the temperature gradient, *W* ∈ [65,320], ensuring that their thermal optimum would likely exist after climate change and that there was competitive pressure on both the trailing and leading edges. We simulated each combination of target species type, fraction relocated *ρ* from 0 to 1 (by 0.05), and thermal optimum estimate while keeping consistent values for AM survival probability µ = 0.8, cooldown time *α* = 5, and relocation adjustment *β* = 10. From baseline simulations without AM (Appendix S2), we chose low-population threshold for AM *η* = 42, high enough that relocation could occur before extinction but low enough to avoid relocating species that would have persisted even without AM.

To determine what types of species and communities are conducive to AM success (increased persistence likelihood with AM), we ran random forest classifications (`randomForest` 4.6-14 package, R Version 3.5.1) separately for low and high stochasticity (note the analogous analysis for vulnerability to climate change in simulations without AM in Appendix S2). The dependent variable was the fate of the target species: global extinction or persistence throughout all locations, disregarding the fate in the original range. The independent variables were target species’ thermal optimum (*z*_*F*_), difference in thermal optimum between target species and neighbors (*z*_diff,*P*_, *z*_diff,*E*_), target and neighbor species’ dispersal (*γ*_*F*_, *γ*_*P*_, *γ*_*E*_), target and neighbor species’ thermal tolerance breadths (*σ*_*F*_, *σ*_*P*_, *σ*_*E*_); target and neighbor species’ initial population sizes (*N*_*F*_(0), *N*_*P*_(0), *N*_*E*_(0)), inverse Simpson’s diversity index of the initial community (*D*_*W*_), measured temperature change (*c*_*T*_), measured standard deviation in temperature (*s*_*T*_), and the deviation between the estimated and true thermal optimum (*z*_est,dev_). To focus on cases where target species specifically benefited from AM, we only included simulations in which the target species went extinct without AM. To simplify analysis, we focused random forest classifications on simulations with fraction-moved *ρ* = 0.5, shortest dispersers, and realized niche estimates.

## Results

Under all scenarios, relocated species had a higher chance of persisting when relocating an intermediate fraction of the total population during AM (Fig. 2, 3a-b). Moreover, target species’ persistence was typically lower when relocating 100% of the total population than without relocation (except when relocating the shortest disperser). Relocating an intermediate fraction of the total population usually allowed the target species to exist in higher densities relative to interacting species in both the donor and recipient communities (Fig. 3c-d). On average, this increased the likelihood of long-term persistence of both the donor and recipient populations (Fig. 3e-f) and overall persistence compared to no management action (Fig. 2, solid/dashed vs. thin dotted lines). Often, AM involved multiple relocations (Fig. 4a-b), and the higher AM success when relocating intermediate fractions came with more individual relocation events (Fig. 4c-d). Assisted migration had little effect on the persistence of non-target species and final community diversity (Appendix S5), so the remaining results focus on persistence instead of invasion risk arising from the competitive interactions modeled here.

**Figure 2:**
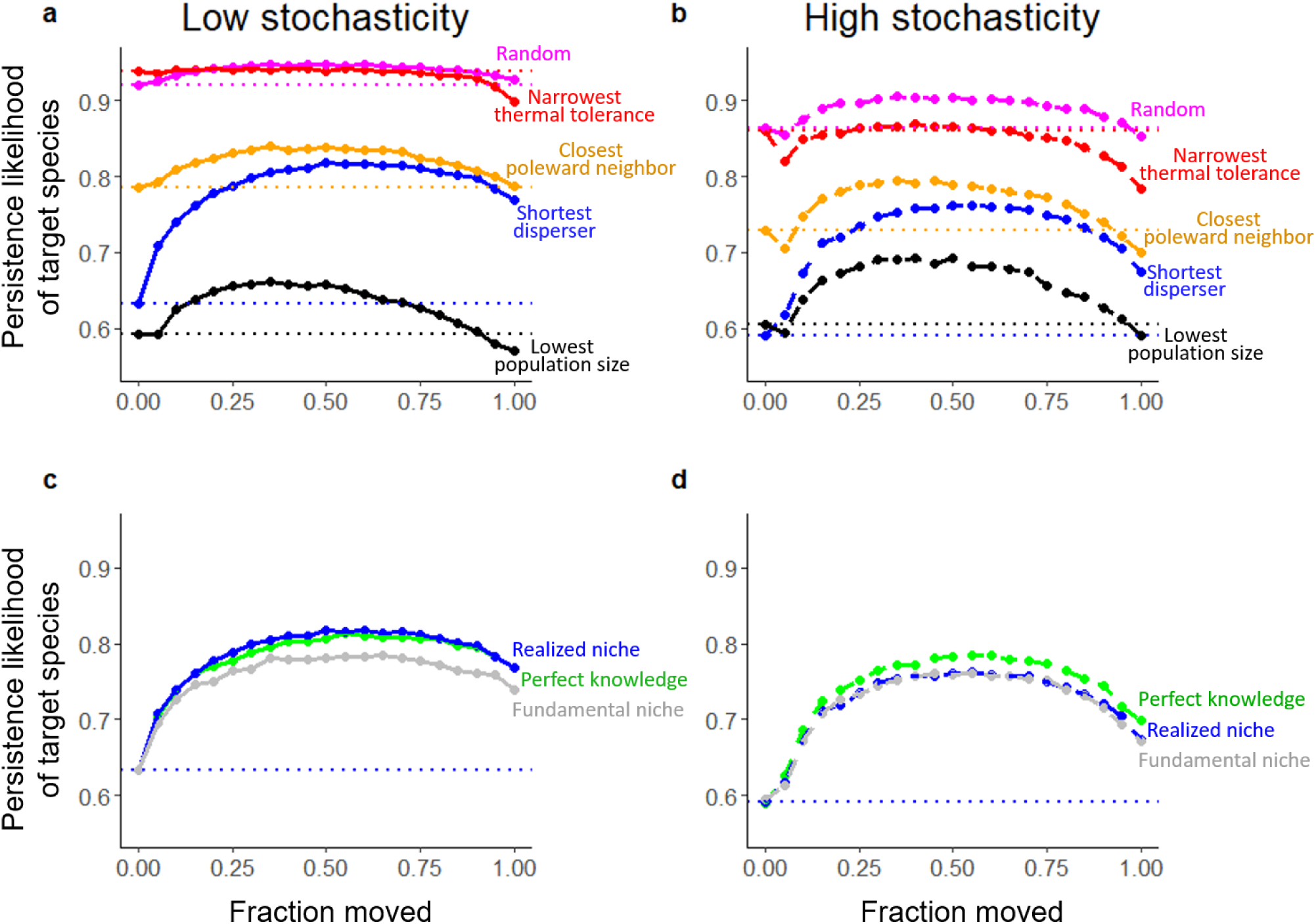
During climate change simulations, the persistence likelihood of a target species chosen for assisted migration (vertical axis) depended on the fraction of that population that was relocated (horizontal axis), and the level of environmental stochasticity (a,c: low, b,d: high). The dotted horizontal lines correspond to persistence likelihood without AM (zero moved, or no management action) and are colored to match each comparison. (a,b) The effect of assisted migration on target species’ persistence with different types of target species chosen for relocation: random, narrow thermal tolerance, closest poleward neighbor, shortest dispersal, or lowest population size. The thermal optimum estimate used in each of these was the realized niche estimates (based on the species initial distribution). (c,d) The effect of assisted migration on target species’ persistence with different types of thermal optimum estimates for relocation decisions: realized niche, fundamental niche, or perfect knowledge. The target species in each of these simulations was the species with the shortest dispersal. Note that the persistence likelihood for non-target species is unaffected (Appendix S5).

**Figure 3:**
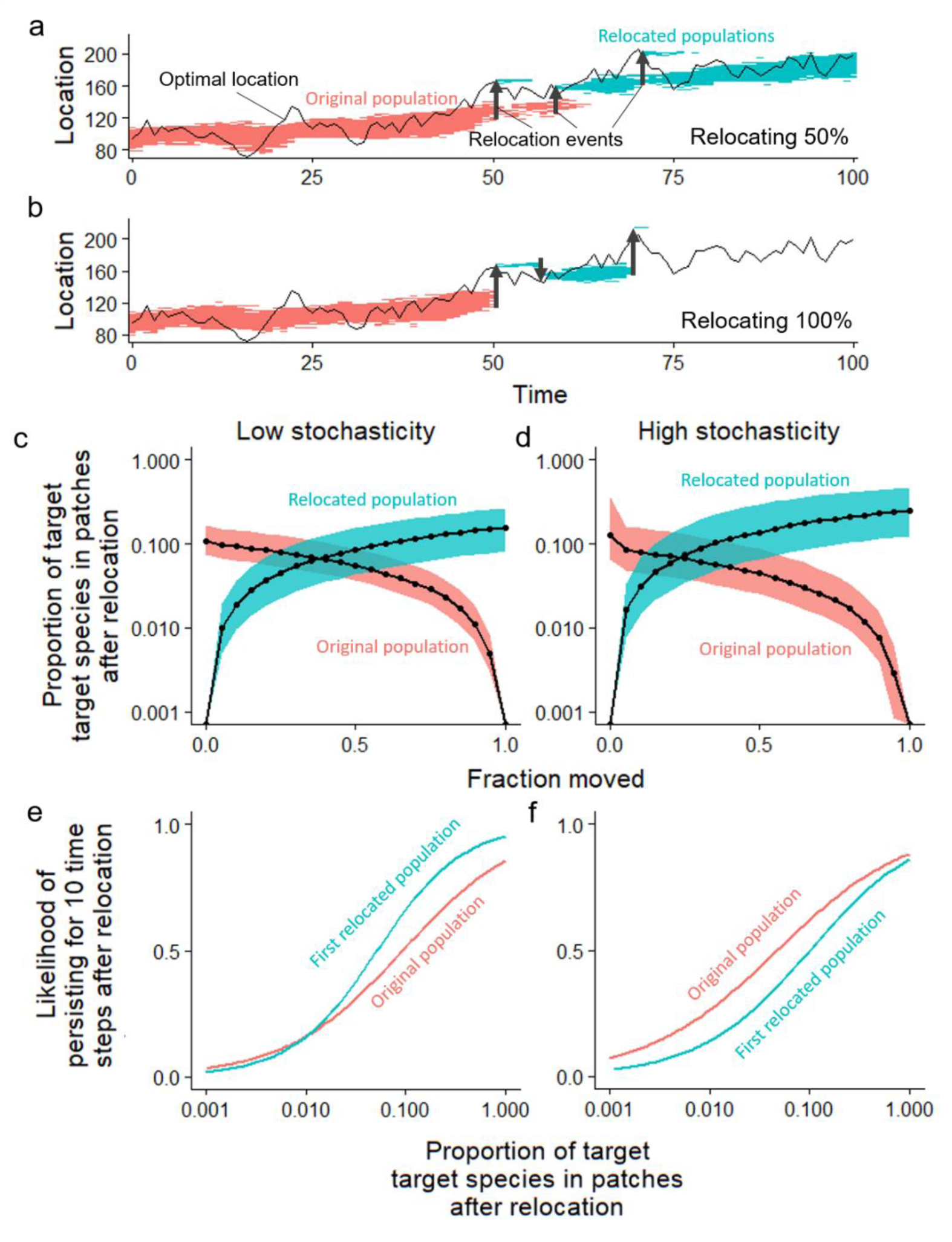
The dynamics and fate of original and relocated populations depend on what fraction of the total population is relocated. (a-b) Example simulations showing how the shortest dispersing species’ range changes over time in a high stochasticity environment. The population follows its optimal climate (thin solid line) as it moves poleward (higher location number). If the total population size falls below a low threshold *η*, a fraction is relocated (thick arrows) based on the estimated thermal optimum. Original and relocated populations are colored separately. (c-d) After relocation, the composition of the donor and recipient communities depend on what fraction of the target species was relocated (horizontal axis). Solid lines represent the median value for the fraction of propagules in the community that are the target species, while the surrounding ribbons represent 25% and 75% quantiles. (e-f) The donor and recipient populations are more likely to persist for at least 10 time steps after relocation (vertical axis) if the target species makes up a higher fraction of the surrounding community (horizontal axis). Lines correspond to predictions from logistic regression.

**Figure 4:**
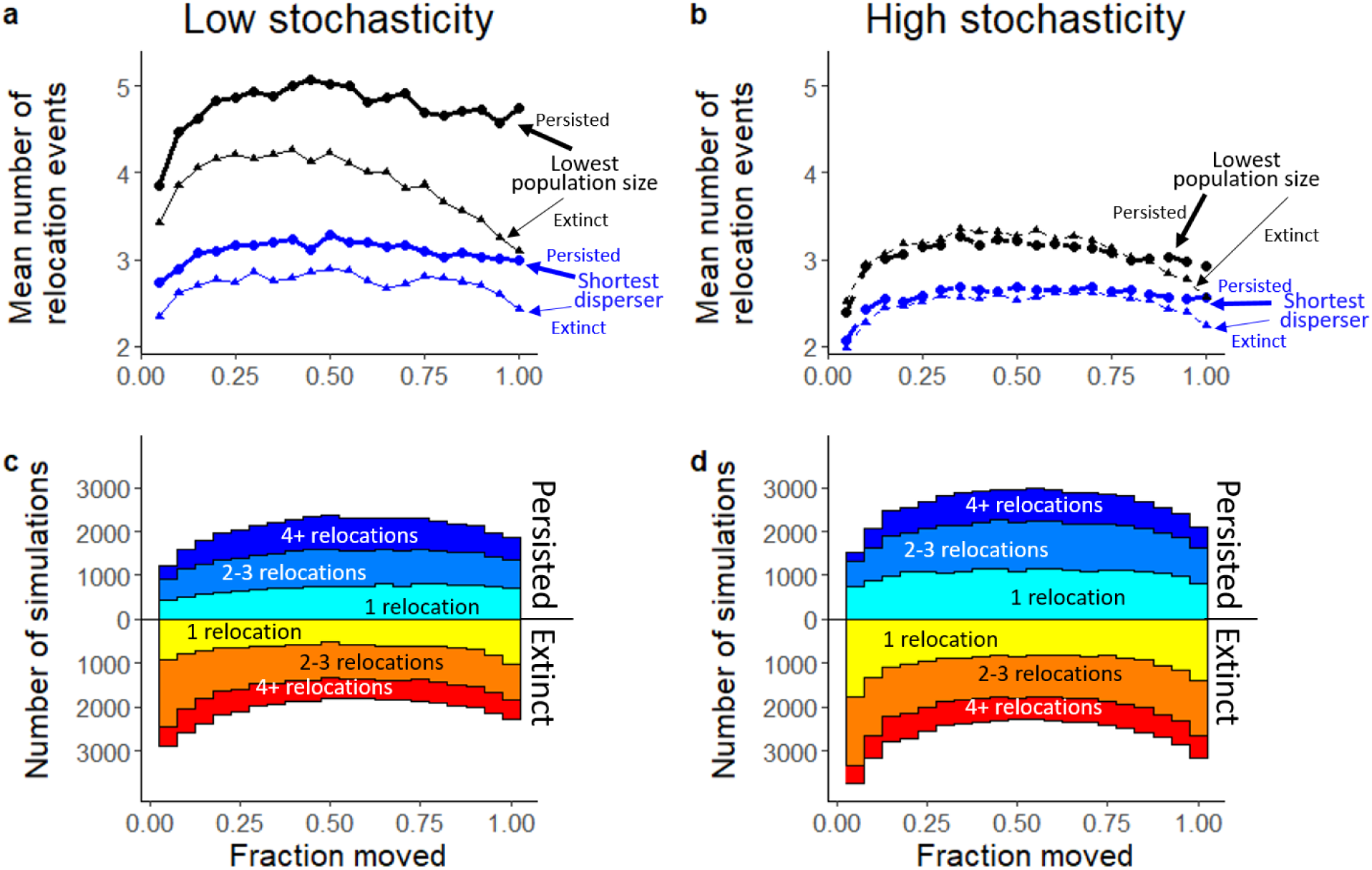
The number of independent relocation events that occurred during assisted migration simulations for low stochasticity (a,c) and high stochasticity (b,d) environments. (a,b) Among species that were relocated at least one time, the mean number of independent relocation events (vertical axis) that occurred under a range of values for the fraction of the population that was relocated each time (horizontal axis). Simulations are separated by target species (color) and whether or not the species went extinct (thick: persisted, thin: extinct). The simulations shown here use the initial distribution (realized niche) estimate of the species’ thermal optimum. (c,d) We categorized assisted migration simulations by fate of the species (persistence/extinction) and the number of independent relocation events that occurred over the course of the simulation. Here we show the number of simulations that fall into these categories (colored differently) depending on the fraction of the population moved during assisted migration (horizontal axis). The center line in each sub-figure separates simulations where the target species survives (below the line) and where it goes extinct (above the line). Simulations shown here use the shortest disperser as the target species and the initial distribution (realized niche) estimate of the species’ thermal optimum.

Of the possible target species, the shortest dispersers experienced the greatest benefit from AM (Fig. 2a-b). For most treatments, AM also increased persistence of target species with the lowest population sizes, species with the closest poleward neighbors, and randomly picked species. However, AM usually decreased persistence of species with the narrowest thermal tolerances (specialists).

On average, AM had a similar effect on persistence regardless of how we estimated the species’ thermal optimum (Fig. 2c-d). Under high stochasticity, AM was most successful with perfect knowledge of species’ thermal optima, but under low stochasticity, AM was most successful with realized niche estimates. This difference suggests stronger competition in low-stochasticity environments such that competition set species limit more than species’ inherent thermal tolerances.

For both levels of stochasticity, three of the top four most important variables for predicting AM success of the shortest disperser were the target species’ initial population size *N*_*F*_(0), the target species’ thermal tolerance breadth *σ*_*F*_, and the difference in thermal optimum between the target species and its poleward neighbor *z*_diff,*P*_ (random forest classifications; out-of-bag error: 25.27% low stochasticity, 30.56% high stochasticity) (Fig. 5a-b). AM was most successful when the values of these characteristics were higher (Fig. 5c-e), suggesting that AM is most likely to benefit generalists with higher population sizes and less poleward competition. Under low stochasticity, AM was less successful if the poleward neighbor was a specialist with narrow thermal tolerance breadth *σ*_*P*_ (Fig. 5f), implying that poleward competition limited AM success under low stochasticity but not as much under high stochasticity. AM was also more successful when thermal optimum estimates were warmer than the true value (positive deviation of *z*_est,dev_) (Fig. 5g), and this effect was stronger under high stochasticity. Colder estimates placed target species further along the climate gradient, often beyond temperatures under which they can survive, so extreme year-to-year temperature change under high stochasticity would be more likely to drive the relocated population extinct if they are placed into the wrong location.

**Figure 5:**
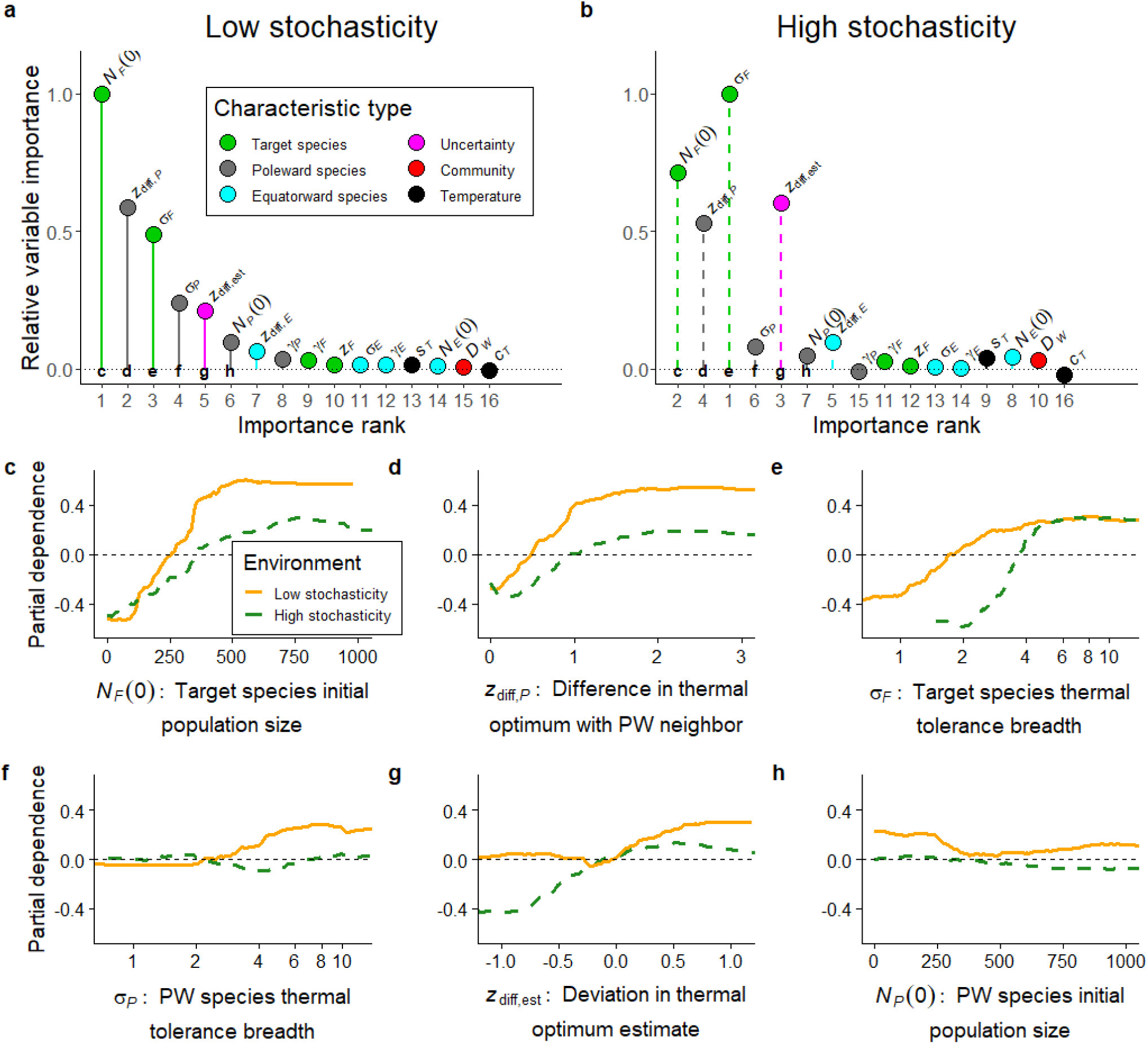
Importance of ecological characteristics from random forest classifications in determining whether assisted migration was successful for the target species (relocating 50% of the shortest dispersing species population with a realized niche estimate thermal optimum). (a,b) Relative unscaled permutation importance of independent variables of whether or not assisted migration improved species’ persistence under low stochasticity (a) and high stochasticity (b). The horizontal axis shows the rank of the variable importance compared to other variables (1 being the most important), arranged in the same order for both plots. Each characteristic is colored depending on whether it is a characteristic of the target species, its neighbors, the full community, the environment, or uncertainty around the thermal optimum estimate. (c-h) Partial dependence of the values of 6 independent variables (corresponding to the top 6 important variables for low stochasticity labeled in panel (a) and (b)) on whether or not assisted migration will increase the shortest disperser’s persistence. The vertical axis is the log-odds of whether assisted migration increases persistence of a species (higher being more likely to persist). Solid lines represent the low stochasticity environment and dashed lines represent the high stochasticity environment. See Table 1 for definitions of symbols.

## Discussion

Our model suggests that assisted migration (AM) can be effective at increasing species’ persistence under climate change if the species is limited by short dispersal, small population sizes, and competition (Fig. 2). However, like many related conservation translocations (Fischer & Lindenmayer 2000), we found that AM often fails when the relocated population does not establish, especially for narrow thermal tolerance species. Relocating an intermediate fraction of the population was consistently an optimal strategy (Fig. 2) due to multiple possible drivers, such as allowing additional relocation attempts after establishment failure (Fig. 4) and reducing the degree of negative density dependence in both the donor and recipient populations (Fig. 3).

### Choosing species for AM

Species are vulnerable to climate change for a variety of reasons, ranging from dispersal-limitation (Pearson 2006), to narrow thermal tolerance (Tewksbury et al. 2008), to competitive interactions (Urban et al. 2012). Our model suggests that the effectiveness of AM as a management strategy depends on the driver of vulnerability. In particular, AM might not be appropriate for specialist, narrow-tolerance species. With only a narrow range of temperatures under which specialists can replace themselves, any error in identifying a species’ optimal climate would disproportionately affect specialists. However, because of our assumption of a jack-of-all-trades-master-of-none trade-off (Levins 1968), specialist species usually persisted under climate change without AM (Appendix S2). With this trade-off, specialists were stronger competitors under lottery competition (Chesson & Warner 1981), outweighing the costs of narrow thermal tolerance in our model. Removing this trade-off would reduce specialists’ competitiveness, making them potentially more vulnerable to climate change, but also potentially more vulnerable to having smaller, divided populations with AM.

AM was most successful for dispersal-limited species because, in these cases, AM directly mitigated the driver of their vulnerability to climate change (McLachlan et al. 2007; Schwartz et al. 2012). Not only was short dispersal a strong predictor of extinction (Appendix S2), but the shortest dispersers also had the strongest proportional increase in persistence with AM (Fig. 2). Moreover, the shortest dispersers were the only target species with increased average persistence in every variation of AM that we modeled. We also found that species with low initial population sizes and species with close poleward competitors were also likely to be vulnerable to climate change and to benefit from AM (Fig. 2). Though these species also had lower-than-average dispersal (Appendix S6), they were likely to be strong AM candidates because of their other vulnerabilities (Thompson & Gonzalez 2017). For example, species with low population sizes might disperse far but not produce enough offspring to realize the full extent of their dispersal potential. Given the possibility that climate change could select for increased dispersal, fecundity, and climate tolerance in relatively short timeframes (Nadeau & Urban 2019), the traits associated with AM success could change over time. This rapid evolution might act as another source of uncertainty that can affect AM decisions, such as the choice of source locations or whether declining populations reliably indicate the persistence risk (as the “evolutionary rescue” process of adapting to novel environments often entails a period of population decline before eventual adaptation and growth; Carlson et al 2014). Though close poleward competitors also increased extinction risk, our focus on lottery competition (amongst many possible drivers of competitive coexistence; Chesson 2000) ignores the niche differentiation for environmental characteristics beyond temperature, which might reduce the impact of poleward competitors. While acknowledging these assumptions inherent to our simple model, our results suggest that AM might be considered for conserving a variety of species beyond those that are directly dispersal limited.

Even under optimal conditions, AM did not prevent the extinction of nearly 20% of short-dispersing species (Fig. 2). For specialists, AM failed because suitable environments were sparse and narrow, but for other species, AM failed because they had a combination of characteristics that limited establishment success (such as a species with both short dispersal and narrow thermal tolerance) (Fig. 5). In these cases, managers might combine alternative management strategies, like increasing connectivity, removing barriers, or creating new reserves (Heller & Zavaleta 2009; Lawler & Olden 2011). For example, we found that species persisted more often after being relocated into low-density communities (Fig. 3), suggesting that managers could prepare the recipient ecosystems by controlling the populations of resident species (Godefroid et al. 2011). This approach might limit competitive pressure, reduce the risk of establishment failure, and temporarily increase the realized niche of the relocated species, but it would come with additional risks to resident species. To be able to compare AM to alternative management strategies such as increasing connectivity, protection, and restoration, our next step is extending our model to incorporate variable carrying capacity and a two-dimensional landscape. Preliminary results suggest that variable carrying capacities and two-dimensional landscapes do not affect the qualitative take-homes about AM highlighted here.

### Fractional relocation

We found that AM was most successful when we relocated an intermediate fraction of the total population (typically around 50-60%; Fig. 2), as this was most robust to uncertainty that could cause AM actions to fail. By leaving donor population to persist temporarily, this approach retains a source for future conservation actions in case relocation occurs at the wrong time or into the wrong place (Fig. 3, 4). Fractional relocation also buffers against the risk of falsely identifying a target species for AM, in which case leaving some individuals behind could allow the species to recover those individuals that might be lost during AM. This contrasts with past AM models that assume the optimal strategy is to move the entire population, as the lagging donor population would eventually go extinct without management (McDonald-Madden et al. 2011). Though relocating more individuals should increase the chances that a species establishes (Godefroid et al. 2011; Blackburn et al. 2015), this could have diminishing returns with negative density dependence (Fischer & Lindenmayer 2000). Instead, our simulations suggest that, in some cases, relocating a fraction of a species could create two smaller populations, each with less negative density dependence than a single unmanaged population (Fig. 3). However, creating two smaller populations could increase extinction risk in some situations. Though our model accounted for the extinction risk through demographic and environmental stochasticity (Lande 1998), we did not include additional extinction risks from Allee effects or explicit genetic factors that could cause inbreeding depression (Gilpin & Soulé 1986). Note that this conclusion is robust to different values of threshold population size for movement *η* and carrying capacity *K* that influence the likelihood of engaging in AM and degree of density dependence, respectively (Appendix S3).

Fractional relocation could also be robust to other risks that we did not directly model, such as the risk of invasion beyond that arising from competitive interactions. Though fractional relocation relies largely on multiple translocations, which repeatedly expose the recipient ecosystem to potential invasion events (Kolar & Lodge 2001; Lockwood et al. 2005), intentionally relocating fewer individuals into a well-monitored ecosystem could also make it easier to detect and prevent invasions before they occur. Similarly, smaller releases could be easier to control if funding, planning, or societal opinions change, and reversal is necessary (Haight et al. 2000).

### Community ecology of AM

Our model builds on past research that suggests competition can prevent some species from tracking climate change (Urban et al. 2012). Though species were vulnerable to extinction if poleward species were strong competitors, AM was also less successful when we relocated species into an area occupied by stronger competitors (Fig. 3, 5). The effect of competition was higher in competition-driven, low-stochasticity communities where AM success depended on characteristics of the poleward species than in dispersal-driven, high-stochasticity communities where success depended more on the ability to accurately place species into their optimal climates. This difference suggests that historical climate variability and community assembly could inform management decisions about AM. For example, limiting competitive interactions (Godefroid et al. 2011) might be more effective for species from environments with low historical variability, whereas relocating species into climate refugia (Morelli et al. 2016) might be more effective for species from environments with higher historical variability.

Though we did not find any substantial negative effects of relocated species on the persistence of species in recipient communities (Appendix S5), we made several simplifications to our model that could have limited the capacity for invasion impacts to occur. First, the simple spatial structure of our model assumed a single contiguous community that assembled without distinct barriers, making the AM in our model analogous to intra-continental relocations that are less likely to cause invasions (Mueller & Hellmann 2008). Additionally, for the sake of simplicity, our model considered only intra-guild lottery competition without the wider web of species interactions that would naturally occur. Biological invasions usually involve more complex ecological dynamics, many of which are taxon-specific and difficult to generalize (Kolar & Lodge 2001; Lockwood et al. 2005; Simberloff et al. 2013). Enemy-release effects would occur if relocated species escape antagonistic interactions that limits growth within their original range (Prenter et al. 2004). Also, relocated species might carry pathogens or parasites that spread to other species in recipient community (Simler et al. 2018). A richer set of interactions could also complicate AM success, as relocating a species without a mutualist might limit establishment (Plein et al. 2016). More interactions would also allow the analysis of AM on characteristics of ecosystem structure and function beyond the extreme outcome of extinction analyzed here. Overall, while our simple competitive framework provides a first step toward exploring the uncertainties and community context of AM, a more complete set of interactions will be necessary to understand the full range of outcomes that could follow a relocation event, from establishment failure, to invasion, to the wide-scale restructuring of ecological communities that is already taking place with climate change (Alexander et al. 2015; Thompson & Gonzalez 2017).

## Supporting information

Supplementary methods (Appendix S1-S2), supplementary tables and figures (Appendix S3-S6), and R code (Appendix S7) are available online. The authors are solely responsible for the content and functionality of these materials. Queries (other than absence of the material) should be directed to the corresponding author.

