## Appendices for "Identifying robust strategies for assisted migration in a competitive stochastic metacommunity"

#### Appendix S1: Double geometric distribution

In continuous space models of species dispersal, ecologists often use a Laplace or double exponential distribution kernels (Kot et al. 1996; Neubert & Caswell 2000; Urban et al. 2012). This is largely because a Laplace distribution is leptokurtic, or has "fat-tails" compared to a standard Guassian kernel. These fat tails give some individuals a higher chance of dispersing extremely far from their origin, which may explain how some slow-dispersing species could track rapidly changing climates in the past (Clark et al. 1998). Because these rare long dispersal events could play an important role in how dispersal-limited species could track climate change in our model, we used a discrete-space leptokurtic dispersal kernel.

Just as the Laplace distribution is a "double exponential" distribution, we considered dispersal kernel that would be a "double geometric" distribution. In other words, this kernel resembles a geometric distribution moving away from 0 in both the positive and negative directions. This kernel is defined by one parameter $q_i$, the probability of a propagule remaining in any particular patch (besides the origin). Each propagule either stays at its origin with probability $\frac{q_i}{2-q_i}$ or moves one space in the pole-ward or equator-ward direction, each with equal probability $\frac{1-q_i}{2-q_i}$. Then each propagule either stays at that current patch with probability $q_i$ or continues to move one space in the same direction with probability $1 - q_i$. This process continues until the propagule stays. The probability that a propagule disperses from patch $x$ to path $y$ is then,

$$k_i(x, y) = \left(\frac{q_i}{2 - q_i}\right)(1 - q_i)^{|x-y|}. \tag{1}$$

Using this probability mass function, we can determine the mean dispersal distance of a propagule, $\gamma_i$, as the mean absolute value of a random variable $y \sim k_i(x, y)$. That is

$$
\begin{aligned}
E[Y] &= \sum_{y=-\infty}^{\infty} |x-y|\, k_i(x,y) \\
&= \sum_{y=-\infty}^{\infty} |x-y| \left(\frac{q_i}{2-q_i}\right) (1-q_i)^{|x-y|} \\
&= \left(\frac{1}{2-q_i}\right) \left[ \sum_{y=-\infty}^{x-1} |x-y|\, q_i (1-q_i)^{|x-y|} + 0 + \sum_{y=x+1}^{\infty} |x-y|\, q_i (1-q_i)^{|x-y|} \right] \\
&= 2\left(\frac{1}{2-q_i}\right) \sum_{y=0}^{\infty} y\, q_i (1-q_i)^{y} \\
&= \frac{2(1-q_i)}{q_i(2-q_i)}. \qquad (2)
\end{aligned}
$$

*References*


Clark JS et al. 1998. Reid's paradox of rapid plant migration: Dispersal theory and interpretation of paleoecological records. BioScience **48**:13–24.

Kot M, Lewis MA, van den Driessche P. 1996. Dispersal data and the spread of invading organisms. Ecology **77**:2027–2042.

Neubert MG, Caswell H. 2000. Demography and dispersal: Calculation and sensitivity analysis of invasion speed for structured populations. Ecology **81**:1613–1628.

Urban MC, Tewksbury JS, Sheldon KS. 2012. On a collision course: Competition and dispersal differences create no-analogue communities and cause extinctions during climate change. Proceedings of the Royal Society B **279**:2072–2080

#### Appendix S2: Baseline simulations with no assisted migration

We modeled metacommunity dynamics under climate change without AM to determine which characteristics related to species, communities, and environments could predict species' vulnerability when unmanaged. After generating $2^{16}$ initialized communities under both low and high stochasticity, we iterated through 100 time steps with mean annual temperature change $\tau = 0.04$°C/year to reflect projected temperature changes under RCP8.5 (Urban et al. 2012, IPCC 2014). From these no-AM simulations, we chose the low-population threshold for AM $\eta = 42$, high enough that relocation could occur before extinction but low enough to avoid relocating species that would have persisted even without AM (Appendix S1).

To determine which ecological characteristics could best predict species' vulnerability to climate change, we ran random forest classifications (`randomForest` 4.6-14 package, R Version 3.5.1) on all simulations without AM (separately for low and high stochasticity). The dependent variable was the fate of a single random species after climate change: global extinction or persistence throughout all locations, disregarding the fate in the original range. The independent variables were target species' thermal optimum ($z_F$), difference in thermal optimum between target species and neighbors ($z_{\text{diff},P}$, $z_{\text{diff},E}$), target and neighbor species' dispersal ($\gamma_F$, $\gamma_P$, $\gamma_E$), target and neighbor species' thermal tolerance breadths ($\sigma_F$, $\sigma_P$, $\sigma_E$); target and neighbor species' initial population sizes ($N_F(0)$, $N_P(0)$, $N_E(0)$), inverse Simpson's diversity index of the initial community ($D_W$), measured temperature change ($c_T$), and measured standard deviation in temperature ($s_T$). Because persistence was more common than extinction, we down-sampled for equal sample sizes. The unscaled permutation variable importance of each independent variable estimated how well these characteristics predicted vulnerability and partial dependence quantified the marginal effect of the characteristics on vulnerability.

Without assisted migration (AM), 91.3% of species persisted under climate change in low-stochasticity environments compared with 84.7% persistence under high stochasticity (Fig. S2.1). In both cases, persistence depended most strongly on a small number of characteristics (random forest classifications, out-of bag error: 8.09% low stochasticity, 11.27% high stochasticity; Fig. S2.2). Persistence was lowest when a species had low initial population sizes $N_F(0)$, short dispersal distances $\gamma_F$, and a close poleward neighbors $z_{\mathrm{diff},P}$. Under low stochasticity, persistence depended on the thermal tolerance of the poleward neighbor $\sigma_P$, such that specialists (species with narrow thermal tolerance breadth) with specialists on their leading edge were less likely to persist than specialists with generalists on their leading edge (Fig S2.3). Comparatively, dispersal and the measured standard deviation in temperature ($s_T$) were more important for persistence under high stochasticity. Altogether, competition largely determined persistence under low stochasticity, whereas dispersal largely determined persistence under high stochasticity.

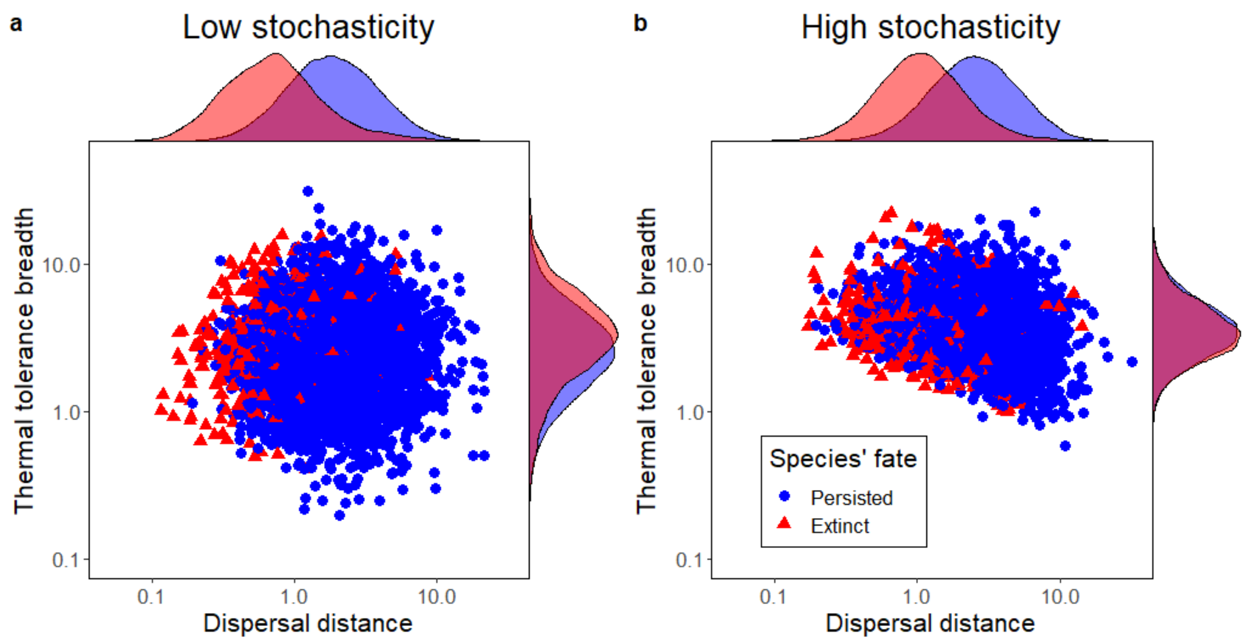


**Figure S2.1**: Species' persistence and the characteristics that predicted persistence varied with environmental stochasticity. Each point shows the fate of a single species (triangle: extinction; circle: persistence) following climate change from a subset of unique simulations plotted over dispersal distance (horizontal axis) and thermal tolerance breadth (vertical axis). On the top and right of these plots are the marginal distributions of these parameters, separated by species' fate. Each axis is on a logarithmic scale.

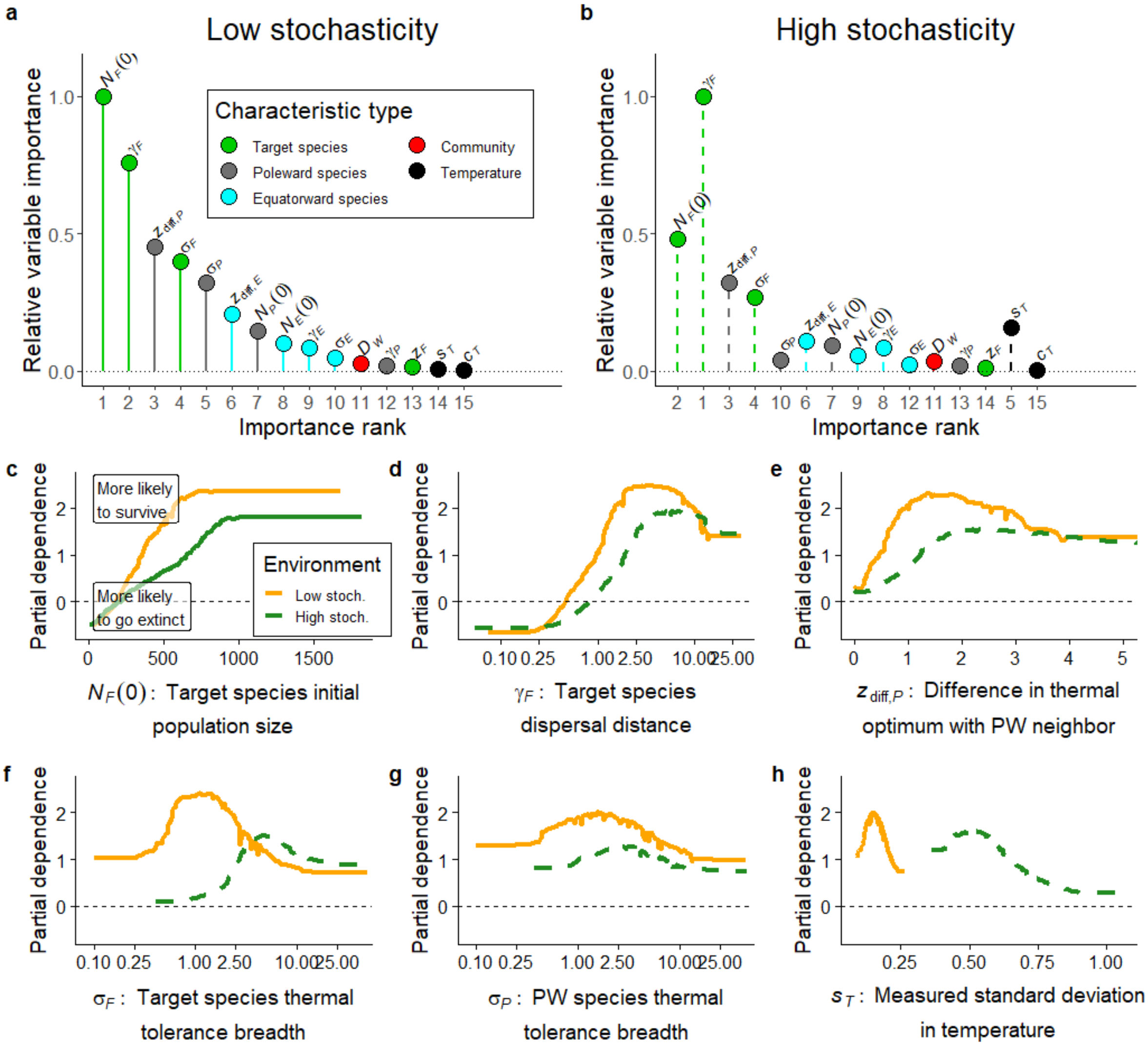


**Figure S2.2**: Importance of species, community, and environmental characteristics in determining whether the target species will persist with climate change without assisted migration. (a,b) Relative unscaled permutation importance of independent variables of whether a species persisted through climate change with no management under low stochasticity (a) and high stochasticity (b). The horizontal axis shows the rank of the variable importance compared to other variables (1 being the most important), arranged in the same order for both plots. Each characteristic is shaded depending on whether it is a characteristic of the target species, its neighbors, the full community, the environment, or uncertainty around the thermal optimum estimate. (c-h) Partial dependence of the values of 6 independent variables (corresponding to the top 6 important variables for low stochasticity labeled in panel (a) and (b)) on whether or not assisted migration will increase the shortest disperser's persistence. The vertical axis is the log-odds of whether a species persisted (higher being more likely to persist). Solid lines represent the low stochasticity environment and dashed lines represent the high stochasticity environment. See Table 1 (main text) for definitions of symbols.

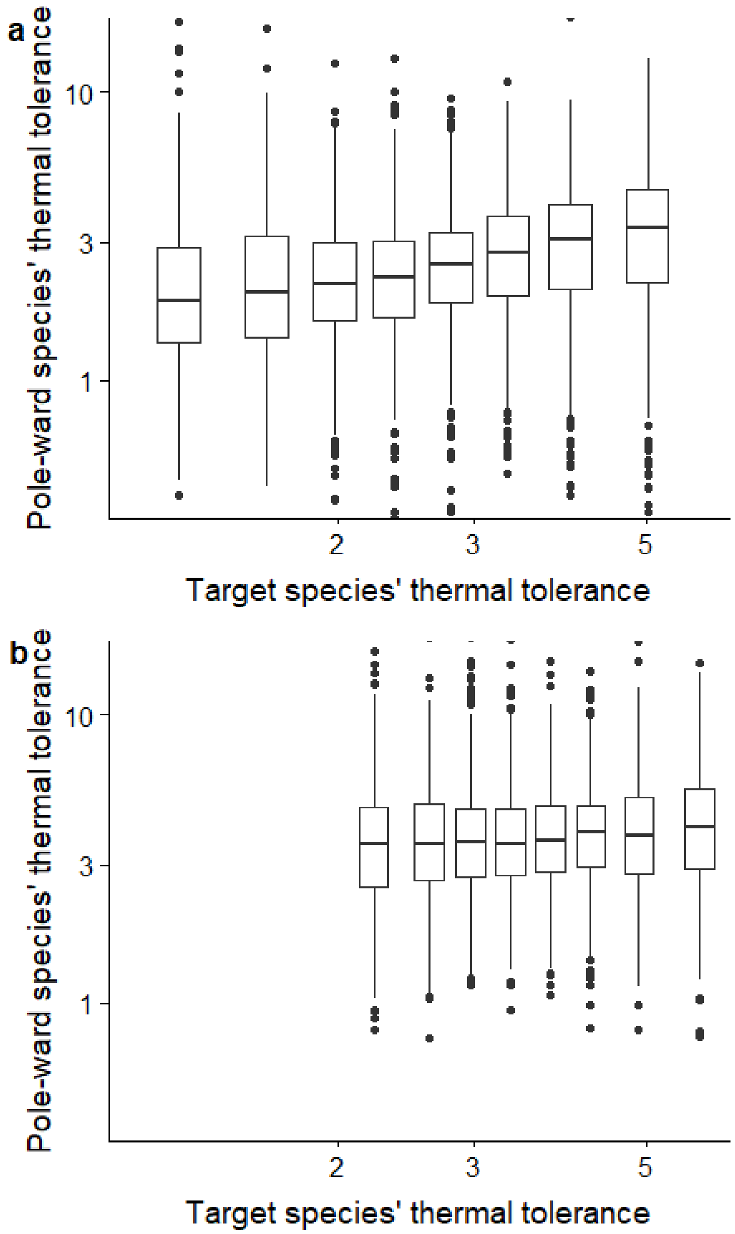


**Figure S2.3**: Comparison of the thermal tolerances of target that went extinct during simulations without assisted migration and with the species on their pole-ward side (a) under low stochasticity and (b) under high stochasticity. Target species are grouped into 10% quantiles of their thermal tolerances with the bottom and top quantiles removed to limit the scale of the figure. Among these species that went extinct, those with higher thermal tolerances had pole-ward neighbors with higher thermal tolerance, with a stronger relationship in lower stochasticity environments.

As we implemented a reactive approach to assisted migration, we had to determine a threshold population below which the population would be relocated. Deciding on appropriate thresholds required us to weigh certain management priorities. If these thresholds were too low, there would likely be only a narrow time window in which managers could react to population decline and relocate the population before extinction. Alternatively, if thresholds are too high, managers might relocate species that are temporarily in decline but not at risk of extinction, risking management funds and effort while potentially creating some extinction risk during an inappropriate relocation.

To find a threshold that balanced these priorities, we analyzed time series of population sizes in the $2^{16}$ high stochasticity simulations in which no management actions were taken. We considered a range of potential population size thresholds from 1 to 100. For each species, if the population fell below the threshold, we determined whether or not the population went extinct following the first instance it fell below the threshold. Those that fell below this threshold but did not go extinct were false positives. Those that fell below the threshold and went extinct in less than 5 time steps were true positives, but impractical to relocate before extinction. Those that fell below the threshold and went extinct in more than 5 but less than 10 time steps were true positives that were practical for relocation. The maximum percentage of practical true positives occurred with a threshold of population size of $\eta = 42$ (Fig. S2.4), which we used as the low population threshold throughout our AM simulations.

*References*

Urban MC, Tewksbury JS, Sheldon KS. 2012. On a collision course: Competition and dispersal differences create no-analogue communities and cause extinctions during climate change. Proceedings of the Royal Society B **279**:2072–2080

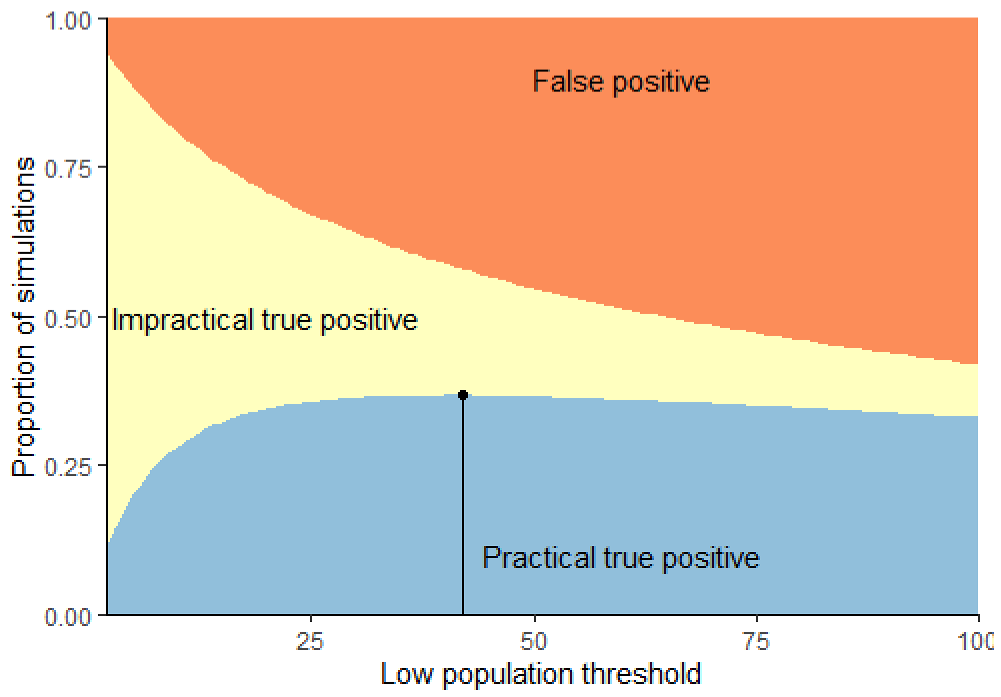


**Figure S2.4**: When simulated under climate change without assisted migration, species fell into three categories (shaded) depending on their fate following the first time they fell below a low population threshold. We compared threshold values to determine when we could detect a species was likely to go extinct, but with enough time to take management action to prevent that extinction. “False positive” species continued to persist until the end of the simulation. “Impractical true positive” species went extinct within 5 time steps. “Practical true positive” species went extinct in more than 5 time steps but less than 10 time steps. The threshold value that optimizes the percentage of practical true positives (42) is marked by the vertical line.

#### Appendix S3: Global sensitivity of $\eta$ and $K$

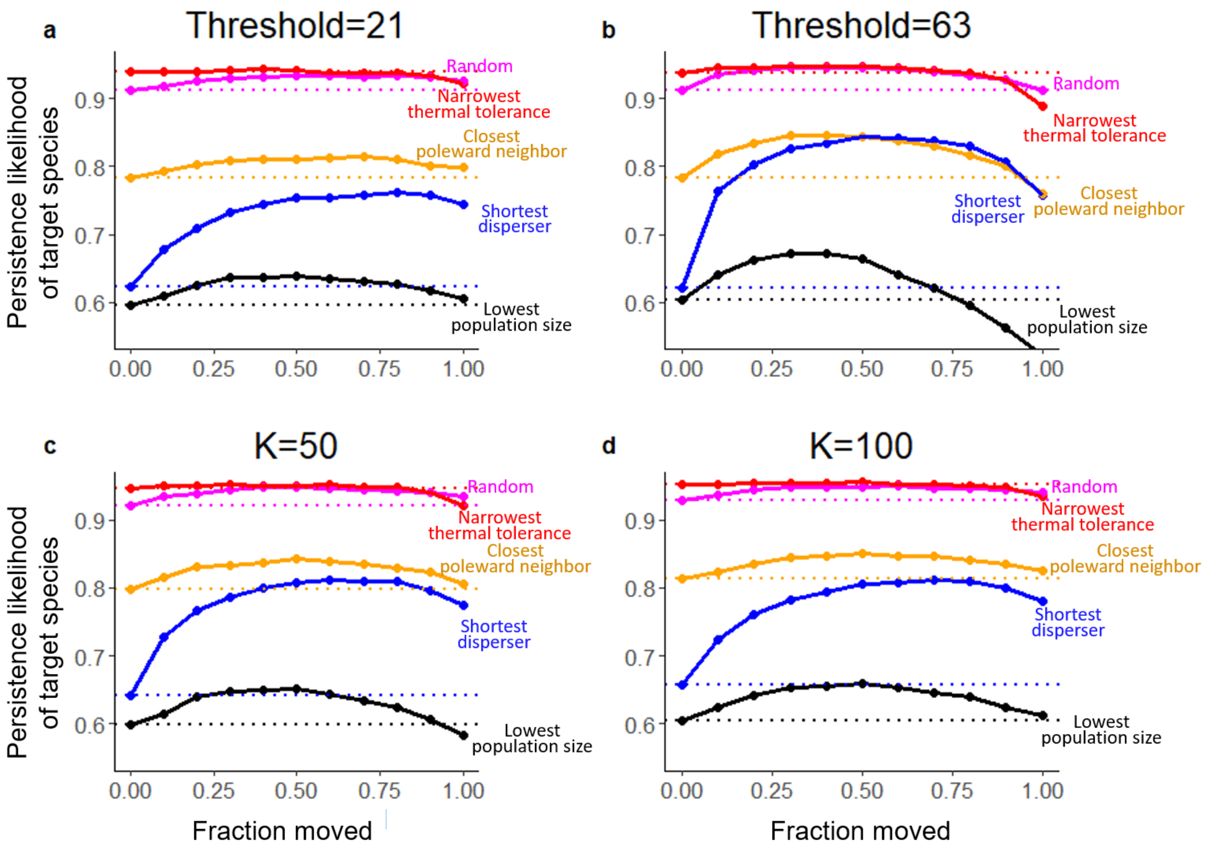


**Figure S3**: Based on 10000 climate change simulations, the persistence likelihood of a target species chosen for assisted migration (vertical axis) depended on the fraction of that population that was relocated (horizontal axis). In this figure, we compare variations on parameter values for the low population threshold for assisted migration ($\eta$) and the community-wide patch carrying capacity ($K$). Comparative default parameter simulations are found in Figure 3 in the main text. In (a) and (b) we use variations on the low population threshold $\eta = 21$ and $\eta = 63$ (compared to a default of $\eta = 42$) while keeping all other parameters at default values and low stochasticity ($\psi = 0.1639$). In (c) and (d), we use variations on the carrying capacity $K = 50$, and $K = 100$ (compared to a default of $K = 30$) while keeping all other parameters at default values. In each set of simulations, the target species chosen for relocation were random, narrow thermal tolerance, closest poleward neighbor, shortest dispersal, or lowest population size. The thermal optimum estimate used in each of these was the realized niche estimates (based on the species initial distribution). The dotted horizontal lines correspond to persistence likelihood without AM (zero moved, or no management action) and are colored to match each comparison.

### Appendix S4: Initialized community data

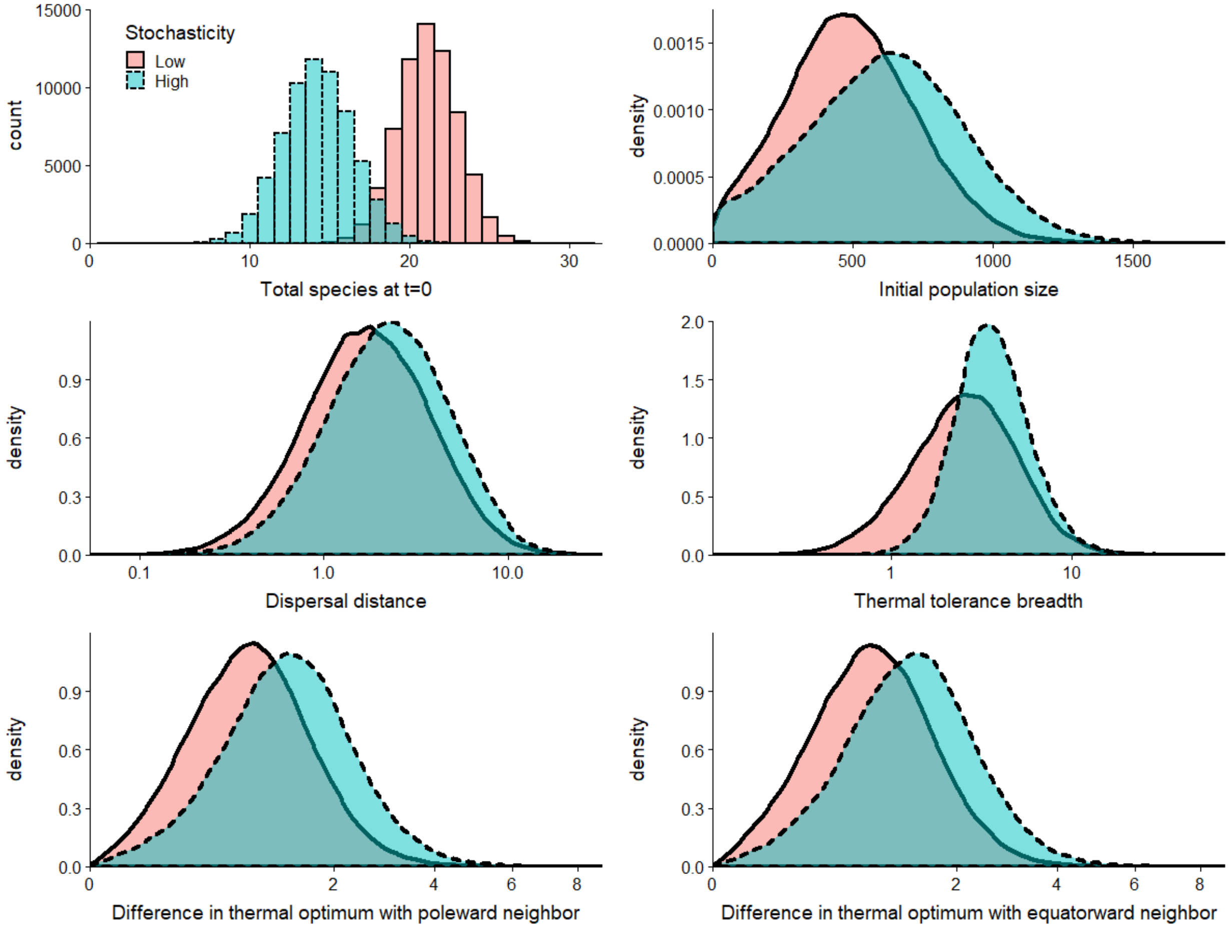


**Figure S4**: Distribution of initialized community characteristics before $2^{16}$ climate change simulations, comparing between low stochasticity (salmon colored with solid line) and high stochasticity (teal colored with dashed line). (a) Number of extant species in the entire spatial gradient (not just the interior sub-region $W$) prior to climate change. (b-f) Characteristics of a species that was randomly picked from those extant in the interior sub-region of the climate gradient $W$ at prior to climate change at time $t = 0$. The horizontal axes are linear in (a-b), log-transformed in (c-d), and square-root-transformed in (e-f).

#### Appendix S5: Effects of assisted migration on non-target species and community

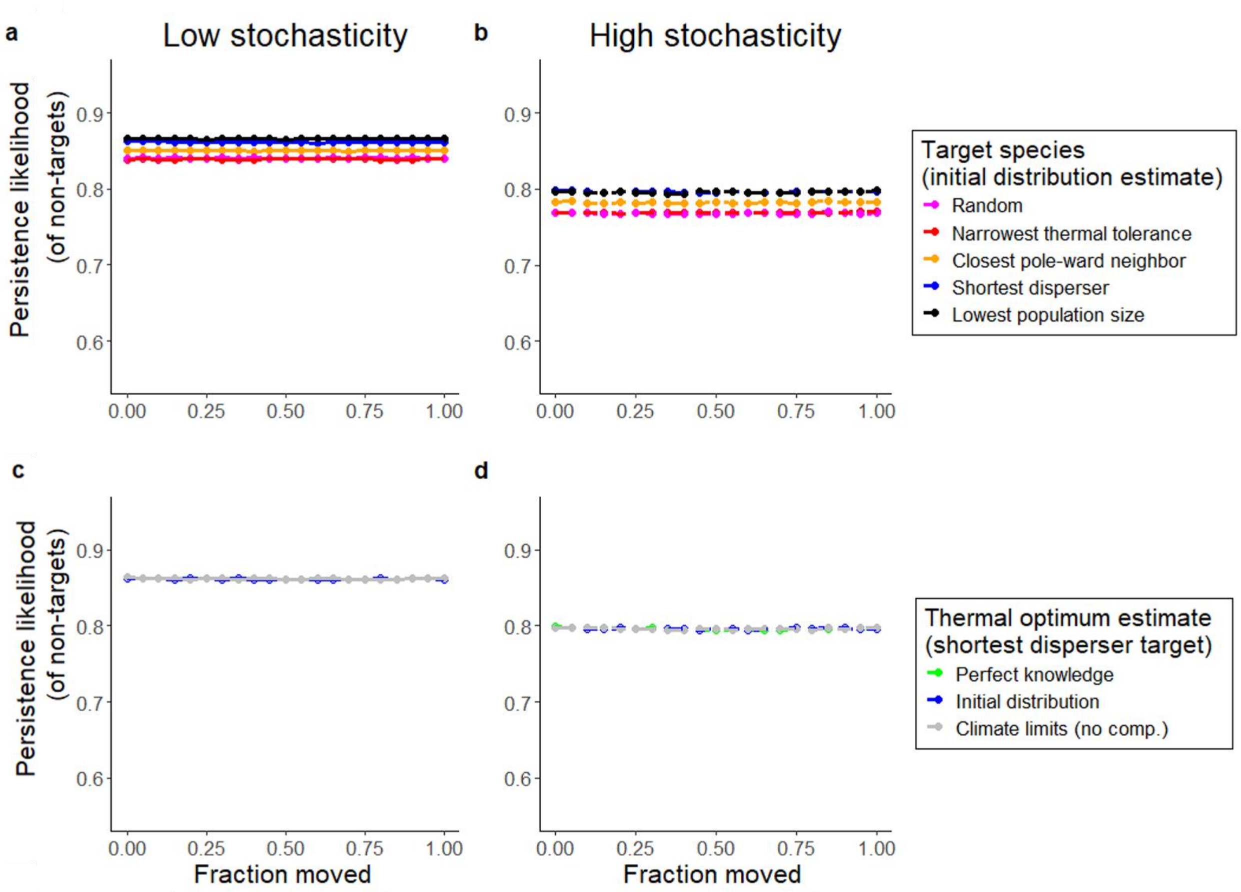


**Figure S4.1**: During climate change simulations, the persistence likelihood of non-target species that were not chosen for assisted migration (vertical axis) did not depend on the fraction of that population that was relocated (horizontal axis). The dotted lines correspond to persistence with no management action and are shaded to match each comparison. (a,b) The effect of assisted migration on non-target species' persistence with different types of target species chosen for relocation. The thermal optimum estimate used in each of these was the realized niche estimates (based on the species initial distribution). (c,d) The effect of assisted migration on non-target species' persistence with different types of thermal optimum estimates. The target species in each of these simulations was the species with the shortest dispersal.

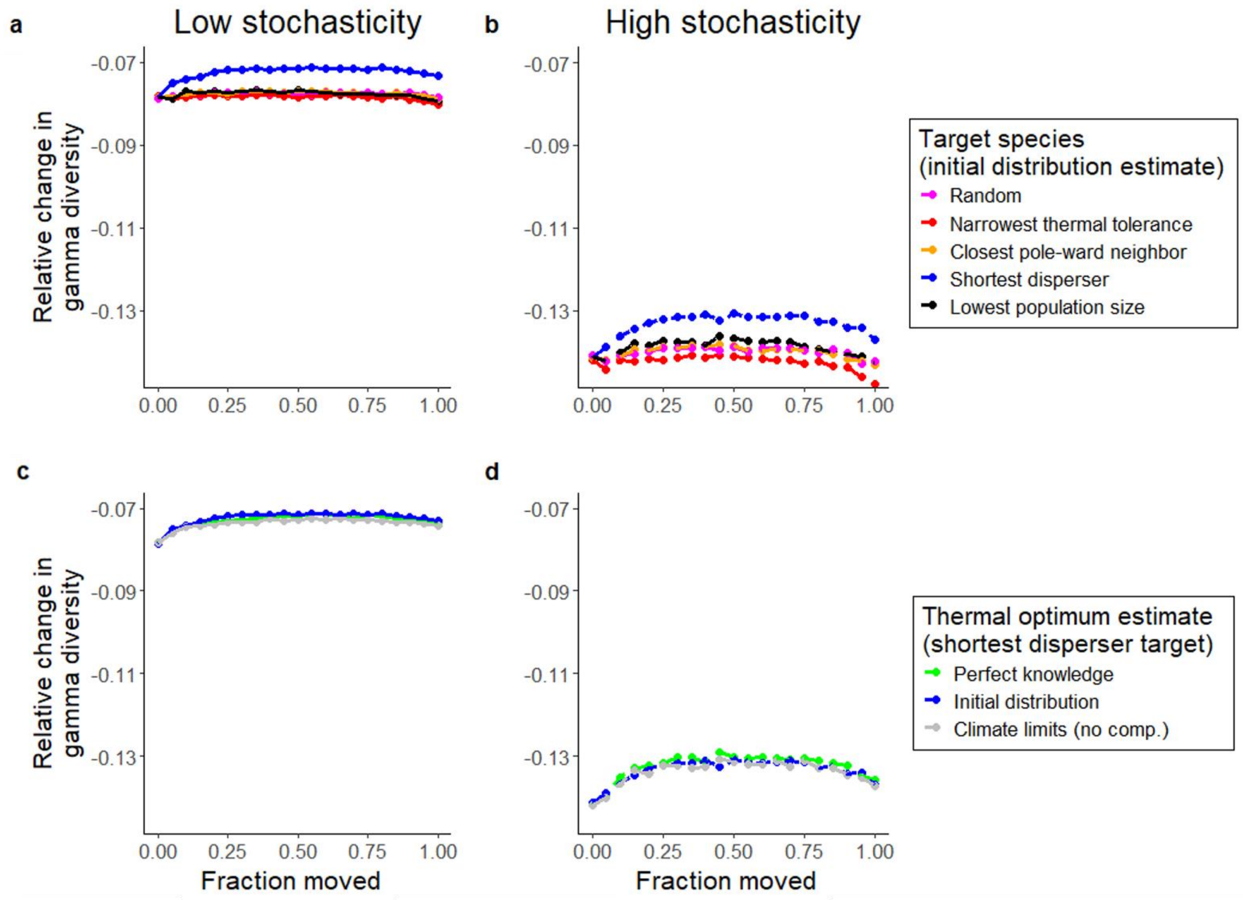


**Figure S4.2**: During climate change simulations, the relative change in gamma inverse Simpson's diversity index (vertical axis) depended on the fraction of the target population that was relocated (horizontal axis). The dotted lines correspond to gamma diversity with no management action and are shaded to match each comparison. Because the persistence likelihood of non-target species is constant across simulations (Fig. S561), the small differences in gamma diversity observed here are likely due to the differences in persistence of the target species. (a,b) The effect of assisted migration on gamma diversity with different types of target species chosen for relocation. The thermal optimum estimate used in each of these was the realized niche estimates (based on the species initial distribution). (c,d) The effect of assisted migration on gamma diversity with different types of thermal optimum estimates. The target species in each of these simulations was the species with the shortest dispersal.

#### Appendix S6: Table of relative values of target species

**Table S5**: Mean value of characteristic values for different types of target species relative to mean values from randomly chosen species in both low and high stochasticity environments.

| Target | Shortest $\gamma_F$ | | Narrowest $\sigma_F$ | | Lowest $N_F(0)$ | | Closest $z_{\text{diff},P}$ | |
|---|---|---|---|---|---|---|---|---|
| Stochasticity | Low | High | Low | High | Low | High | Low | High |
| $z_F$ | 1.00 | 1.01 | 1.00 | 1.00 | 1.00 | 1.01 | 1.00 | 0.99 |
| $\gamma_F$ | 0.22 | 0.27 | 1.06 | 1.48 | 0.85 | 0.83 | 0.90 | 0.97 |
| $\sigma_F$ | 0.98 | 1.10 | 0.28 | 0.47 | 1.49 | 1.24 | 0.92 | 0.99 |
| $N_F(0)$ | 0.85 | 0.84 | 0.88 | 0.99 | 0.29 | 0.39 | 0.73 | 0.77 |
| $z_{\text{diff},P}$ | 0.98 | 1.01 | 0.86 | 0.99 | 0.70 | 0.73 | 0.17 | 0.28 |
| $z_{\text{diff},E}$ | 0.97 | 1.01 | 0.85 | 0.99 | 0.74 | 0.77 | 1.19 | 1.19 |

#### Appendix S7: R code

Sample R code can be accessed at:

https://anonymous.4open.science/r/9e80ce08-9937-4e71-8a77-73e47fac8f15/
